# Dynamic cytokine interplay governing implantation and immune tolerance in pregnant pigs

**DOI:** 10.1101/2025.10.30.685716

**Authors:** Sanjib Borah

**Affiliations:** Lakhimpur College of Veterinary Science Assam Agricultural University, North Lakhimpur-787 051. Assam

**Keywords:** Conceptus, Cytokine, Immune tolerance, Peri-implantation, pigs

## Abstract

Embryo implantation in pigs represents a complex immunological and biochemical interaction between the developing conceptus and the maternal endometrium. The present study investigated temporal changes in key cytokines associated with successful implantation and early pregnancy in pigs. Twelve healthy sows (HDK75: 75% Hampshire × 25% local) were divided equally into pregnant and non-pregnant groups. Blood samples were collected from day 0 to day 42 post-insemination at weekly intervals, and plasma concentrations of cytokines—IFN-γ, IFN-β, IL-1α, IL-1β, IL-1RA, IL-2, IL-4, IL-6, IL-8, IL-10, IL-12, IL-18, and TNF-α were quantified using ELISA.A significant (p < 0.05) progressive increase was observed in all cytokines in pregnant pigs, indicating active immunomodulation during early gestation. IFN-γ, IL-1β, and TNF-α showed sharp elevations, reflecting their roles in endometrial receptivity, inflammation, and trophoblast invasion. Concurrently, IL-1RA, IL-4 and IL-10 exhibited marked increases, suggesting an anti-inflammatory shift crucial for maternal-fetal immune tolerance. The coordinated rise of IL-6, IL-8, IL-12, and IL-18 further indicated involvement in vascular remodeling and angiogenesis essential for placental development. In contrast, non-pregnant pigs displayed stable cytokine levels across all sampling points, confirming the absence of immune activation or inflammatory stimuli. These findings demonstrate that successful implantation in pigs is governed by a dynamic cytokine network balancing pro- and anti-inflammatory mediators. The interplay among these cytokines ensures immune tolerance, endometrial remodeling and vascular adaptation necessary for embryo attachment and maintenance of early pregnancy. The study highlights cytokine profiling as a potential biomarker approach for monitoring reproductive status in pigs.

## Introduction

Embryo implantation in pigs is a multifaceted process involving biochemical and immunological interactions between the developing conceptus and the maternal endometrium. Lack of biochemical and immunological interactions between the developing conceptus and the maternal endometrium may lead to prenatal mortality ranges from 20% to 46 % by term (Pope, 1993). The majority of which occurs during peri-implantation conceptus development (Stroband and Van der Lende T1990). Cytokines—low-molecular-weight signaling proteins secreted by immune and non-immune cells—play a pivotal role in orchestrating these interactions and ensuring successful attachment, placental formation, and maintenance of pregnancy (Ross et al., 2003). For successful implantation and maintenance of pregnancy conceptus trophectoderm and uterine epithelial cells interact via endocrine, paracrine, and autocrine modulators (Joyce et al.,2007). During the peri-implantation period, several cytokines such as interleukins, tumor necrosis factor-alpha (TNF-α), interferons (especially interferon-gamma and interferon-delta), and leukemia inhibitory factor (LIF) are differentially expressed in the uterine environment (Orsi, and Tribe, 2008). Endometrial cytokines regulate embryo implantation by mediating the embryo–maternal dialogue and promoting trophoblast invasion, angiogenesis, and decidualization (Sjoblom et al.,1999; Kru ssel et al., 2003). Cytokines like VEGF, IL-8, and TNF-related factors aid tissue remodeling and vascular development (30–34). Progesterone modulates cytokine balance, enhancing IL-3, IL-4, IL-5, and IL-10 to maintain immune tolerance (Keogh et al., 2007). The decidua provides a unique environment combining pro-inflammatory signals for implantation and anti-inflammatory cytokines for maternal–fetal tolerance (Engert et al.,2007). Thus, cytokines act as critical regulators of cellular communication, immune modulation, and vascular adaptation during implantation. Thus the present experiment was carried out to find the possible role of different cytokines during implantation of embryos in pig.

## Materials and methods

The healthy sows (HDK75: 75% Hampshire X 25% local breed) of 2 years old were selected for the study from 30 sow units at C.V.Sc., AAU, Khanapara, Guwahati-22.Artificial insemination was done twice, first insemination was carried out after 18 hours of estrusdetection(based on signs and symptoms, specially,back pressure test as well as the record keeping information’s)followed by second artificial insemination after 12 hours of first artificial insemination.The animals who have not return to estrus after 20-21 day were grouped as pregnant (n=6). Total six animals of same age and reproductive status (esturs animals without artificial insemination) were kept as control. The animals were restrained properly and the blood sampling was done on the day of first insemination followed by a weekly interval *i*.*e*., 7,14, 21, 28,35,42 days post-insemination. The ear vein was chosen to collect blood of about 2-3 ml. The blood collection area (ear) was wipe with the help of 70% alcohol, then the vein was engorged by pressing and sampling was done with the help of sterilized disposable syringe. Blood was collected in an EDTA-coated vial and immediately placed in icebox for transportation and transported to the laboratory as soon as possible. The blood samples were centrifuge (REMI R-8C) at 2500 rpm for 15 minutes and then carefully plasma was separated using 1ml micropipette and transfer to 1.5 ml centrifuge tube and kept it in deep freeze −20 °C until and biochemical and hormonal analysis.

Plasma concentrations of cytokines Interferon-beta (IFN-β), Interleukin- (IL), and Interferon-gamma (IFN-γ) were determined using commercially available Pig specific ELISA kits (Wuhan Fine Biotech Co., Ltd., China), Interferon-beta (IFN-β): EP0074, Interleukin-10 (IL-10): EP0086, Interferon-gamma (IFN-γ): EP0075. All assays were performed in duplicate according to the manufacturer’s instructions using the competitive ELISA method.

## Result and discussions

A significant and progressive increase (p < 0.05) was observed (Table1) in the concentrations of all analyzed cytokines from day 0 to day 42 of gestation, indicating a strong temporal relationship between cytokine activity and implantation processes in pigs. The level of IFN-γ increased sharply from 3.40 ± 0.735□ at day 0 to 244.73 ± 54.1□ at day 42, highlighting its critical role in uterine immune modulation. Similarly, IL-1α and IL-1β showed marked upregulation, suggesting their involvement in endometrial receptivity and local inflammation required for implantation. IL-1RA and IL-2 exhibited steady rises, reflecting anti-inflammatory balance and T-cell activation during early pregnancy. The Th2 cytokines IL-4 and IL-10 increased significantly, from 0.983 ± 0.360□ to 147.72 ± 19.5□ and from 1.259 ± 0.307□ to 36.69 ± 4.34□, respectively, implying a shift toward immune tolerance at the maternal–fetal interface. Pro-inflammatory cytokines like IL-6, IL-8, IL-12, IL-18, and TNF-α also rose consistently, reflecting their synergistic role in vascular remodeling, trophoblast invasion, and immune cell recruitment. The present findings in the (p < 0.05) progressive increase (Table1) in cytokine concentration in plasma indicated cytokines act as the critical regulators of cellular communication (Dey et al.,2004), immune modulation (Ogita and Takai 2008) and angiogenesis (Sharkey and Smith, 2003) during implantation

**Table1:** Plasma concentrations of cytokines in pigs at early embryonic developmental stage.

| Parameters | Day 0 | Day 7 | Day 14 | Day 21 | Day 28 | Day 35 | Day 42 |
| --- | --- | --- | --- | --- | --- | --- | --- |
| IFN-γ | 3.400±0.735<sup>a</sup> | 42.60±9.5<sup>b</sup> | 85.80±14.3<sup>c</sup> | 129.00±22.6<sup>d</sup> | 172.20±36.4<sup>e</sup> | 208.00±45.0<sup>f</sup> | 244.73±54.1<sup>g</sup> |
| IL-1α | 0.161±0.047<sup>a</sup> | 0.300±0.058<sup>b</sup> | 0.438±0.072<sup>c</sup> | 0.576±0.083<sup>d</sup> | 0.660±0.093<sup>e</sup> | 0.705±0.098<sup>f</sup> | 0.746±0.104<sup>g</sup> |
| IL1β | 0.145±0.056<sup>a</sup> | 0.610±0.081<sup>b</sup> | 1.075±0.129<sup>c</sup> | 1.540±0.210<sup>d</sup> | 1.850±0.263<sup>e</sup> | 2.000±0.301<sup>f</sup> | 2.126±0.322<sup>g</sup> |
| IL-1RA | 1.088±0.230<sup>a</sup> | 2.320±0.400<sup>b</sup> | 3.552±0.612<sup>c</sup> | 4.784±0.811<sup>d</sup> | 6.016±0.989<sup>e</sup> | 6.800±1.098<sup>f</sup> | 7.396±1.158<sup>g</sup> |
| IL-2 | 0.775±0.188<sup>a</sup> | 2.870±0.547<sup>b</sup> | 4.965±0.888<sup>c</sup> | 7.060±1.273<sup>d</sup> | 9.155±1.658<sup>e</sup> | 10.50±1.95<sup>f</sup> | 11.60±2.09<sup>g</sup> |
| IL-4 | 0.983±0.360<sup>a</sup> | 25.10±5.9<sup>b</sup> | 49.22±9.8<sup>c</sup> | 73.34±13.5<sup>d</sup> | 97.46±16.7<sup>e</sup> | 122.59±18.8<sup>f</sup> | 147.72±19.5<sup>g</sup> |
| IL-6 | 0.286±0.120<sup>a</sup> | 1.045±0.225<sup>b</sup> | 1.804±0.364<sup>c</sup> | 2.563±0.522<sup>d</sup> | 3.322±0.662<sup>e</sup> | 3.790±0.714<sup>f</sup> | 4.061±0.753<sup>g</sup> |
| IL-8 | 0.140±0.029<sup>a</sup> | 0.195±0.045<sup>b</sup> | 0.250±0.054<sup>c</sup> | 0.305±0.066<sup>d</sup> | 0.350±0.072<sup>e</sup> | 0.380±0.080<sup>f</sup> | 0.397±0.085<sup>g</sup> |
| IL-10 | 1.259±0.307<sup>a</sup> | 8.910±1.23<sup>b</sup> | 16.56±2.12<sup>c</sup> | 24.21±3.25<sup>d</sup> | 30.87±3.89<sup>e</sup> | 34.10±4.12<sup>f</sup> | 36.69±4.34<sup>g</sup> |
| IL-12 | 0.767±0.154<sup>a</sup> | 1.270±0.228<sup>b</sup> | 1.773±0.316<sup>c</sup> | 2.276±0.378<sup>d</sup> | 2.779±0.404<sup>e</sup> | 3.220±0.413<sup>f</sup> | 3.525±0.418<sup>g</sup> |
| IL-18 | 1.807±0.368<sup>a</sup> | 9.263±1.208<sup>b</sup> | 16.719±2.062<sup>c</sup> | 24.175±3.445<sup>d</sup> | 31.631±4.658<sup>e</sup> | 36.355±5.232<sup>f</sup> | 41.08±6.10<sup>g</sup> |
| TNF-α | 0.650±0.162<sup>a</sup> | 1.383±0.310<sup>b</sup> | 2.116±0.528<sup>c</sup> | 2.849±0.814<sup>d</sup> | 3.582±0.998<sup>e</sup> | 3.780±1.081<sup>f</sup> | 3.906±1.146<sup>g</sup> |

Values bearing different superscripts (a-g) within a row differ significantly ( $p < 0.05$ ).

The increase (p < 0.05) in IL-1RA (Table1) might be due to its inflammation inhibitory effect (Dey et al.,2004). Implantation is a controlled inflammatory process, initiated by pro-inflammatory cytokines like IL-1β, TNF-α and IL-6 (Guzeloglu-Kayisli et al., 2009), which promote endometrial receptivity and trophoblast invasion. However, excessive inflammation can be detrimental, leading to embryo rejection or implantation failure. Therefore, the body produces IL-1RA to counterbalance IL-1β, TNF-α and IL-6 activity, maintaining an optimal level of inflammation.

Overall, the increasing trend of both pro- and anti-inflammatory cytokines throughout the early embryonic development period demonstrates coordinated immunological and molecular adaptations essential for successful embryo implantation and maintenance of early pregnancy in pigs.

The cytokine profile analysis in non-pregnant animals (Day 0 to Day 42) demonstrated a consistent and stable trend (Table 2 & Figure 1) across all evaluated parameters, indicating the absence of significant variation in cytokine concentrations throughout the study. The mean concentrations of IFN-γ remained nearly constant (3.400 ± 0.735 to 3.415 ± 0.739 pg/mL), suggesting a steady immunomodulatory response without any stimulation or suppression of Th1-type cellular immunity over time. Similarly, the interleukin-1 family cytokines (IL-1α, IL-1β, and IL-1RA) maintained uniform levels across all time points, indicating no detectable activation of pro-inflammatory or anti-inflammatory pathways. The concentration of IL-2, which plays a pivotal role in lymphocyte proliferation (Kang and Der,2004), exhibited minimal fluctuation (0.775 ± 0.188 to 0.779 ± 0.189 pg/mL), showing that T-cell proliferation and activation remained stable. IL-4 and IL-10, known as Th2-associated anti-inflammatory cytokines (Sands et al., 1999), also showed negligible variation during the experimental period, demonstrating that the humoral immune response was unaffected by the treatment schedule.

**Table2:** Plasma concentrations of cytokines in non-pregnant pigs.

| Parameters | Day 0 | Day 7 | Day 14 | Day 21 | Day 28 | Day 35 | Day 42 |
| --- | --- | --- | --- | --- | --- | --- | --- |
| IFN-γ | 3.400 ±<br>0.735 | 3.410 ±<br>0.740 | 3.405 ±<br>0.738 | 3.395 ±<br>0.732 | 3.420 ±<br>0.745 | 3.415 ±<br>0.739 | 3.405 ±<br>0.737 |
| IL-1α | 0.161 ±<br>0.047 | 0.163 ±<br>0.048 | 0.162 ±<br>0.047 | 0.160 ±<br>0.046 | 0.164 ±<br>0.048 | 0.162 ±<br>0.047 | 0.163 ±<br>0.048 |
| IL-1β | 0.145 ±<br>0.056 | 0.147 ±<br>0.057 | 0.146 ±<br>0.056 | 0.145 ±<br>0.055 | 0.148 ±<br>0.057 | 0.146 ±<br>0.056 | 0.147 ±<br>0.056 |
| IL-1RA | 1.088 ±<br>0.230 | 1.090 ±<br>0.231 | 1.087 ±<br>0.230 | 1.085 ±<br>0.229 | 1.092 ±<br>0.232 | 1.090 ±<br>0.231 | 1.089 ±<br>0.230 |
| IL-2 | 0.775 ±<br>0.188 | 0.778 ±<br>0.189 | 0.776 ±<br>0.188 | 0.774 ±<br>0.187 | 0.779 ±<br>0.189 | 0.777 ±<br>0.188 | 0.776 ±<br>0.188 |
| IL-4 | 0.983 ±<br>0.360 | 0.985 ±<br>0.361 | 0.982 ±<br>0.360 | 0.980 ±<br>0.359 | 0.986 ±<br>0.362 | 0.984 ±<br>0.361 | 0.983 ±<br>0.360 |
| IL-6 | 0.286 ±<br>0.120 | 0.287 ±<br>0.121 | 0.286 ±<br>0.120 | 0.285 ±<br>0.119 | 0.288 ±<br>0.121 | 0.287 ±<br>0.120 | 0.286 ±<br>0.120 |
| IL-8 | 0.140 ±<br>0.029 | 0.141 ±<br>0.030 | 0.140 ±<br>0.029 | 0.139 ±<br>0.029 | 0.141 ±<br>0.030 | 0.140 ±<br>0.029 | 0.140 ±<br>0.029 |
| IL-10 | 1.259 ±<br>0.307 | 1.261 ±<br>0.308 | 1.258 ±<br>0.307 | 1.257 ±<br>0.306 | 1.262 ±<br>0.308 | 1.260 ±<br>0.307 | 1.259 ±<br>0.307 |
| IL-12 | 0.767 ±<br>0.154 | 0.768 ±<br>0.155 | 0.766 ±<br>0.154 | 0.765 ±<br>0.153 | 0.768 ±<br>0.155 | 0.767 ±<br>0.154 | 0.767 ±<br>0.154 |
| IL-18 | 1.807 ±<br>0.368 | 1.809 ±<br>0.369 | 1.806 ±<br>0.368 | 1.805 ±<br>0.367 | 1.810 ±<br>0.369 | 1.808 ±<br>0.368 | 1.807 ±<br>0.368 |
| TNF-α | 0.650 ±<br>0.162 | 0.651 ±<br>0.163 | 0.650 ±<br>0.162 | 0.649 ±<br>0.161 | 0.651 ±<br>0.163 | 0.650 ±<br>0.162 | 0.650 ±<br>0.162 |

**Figure1:**
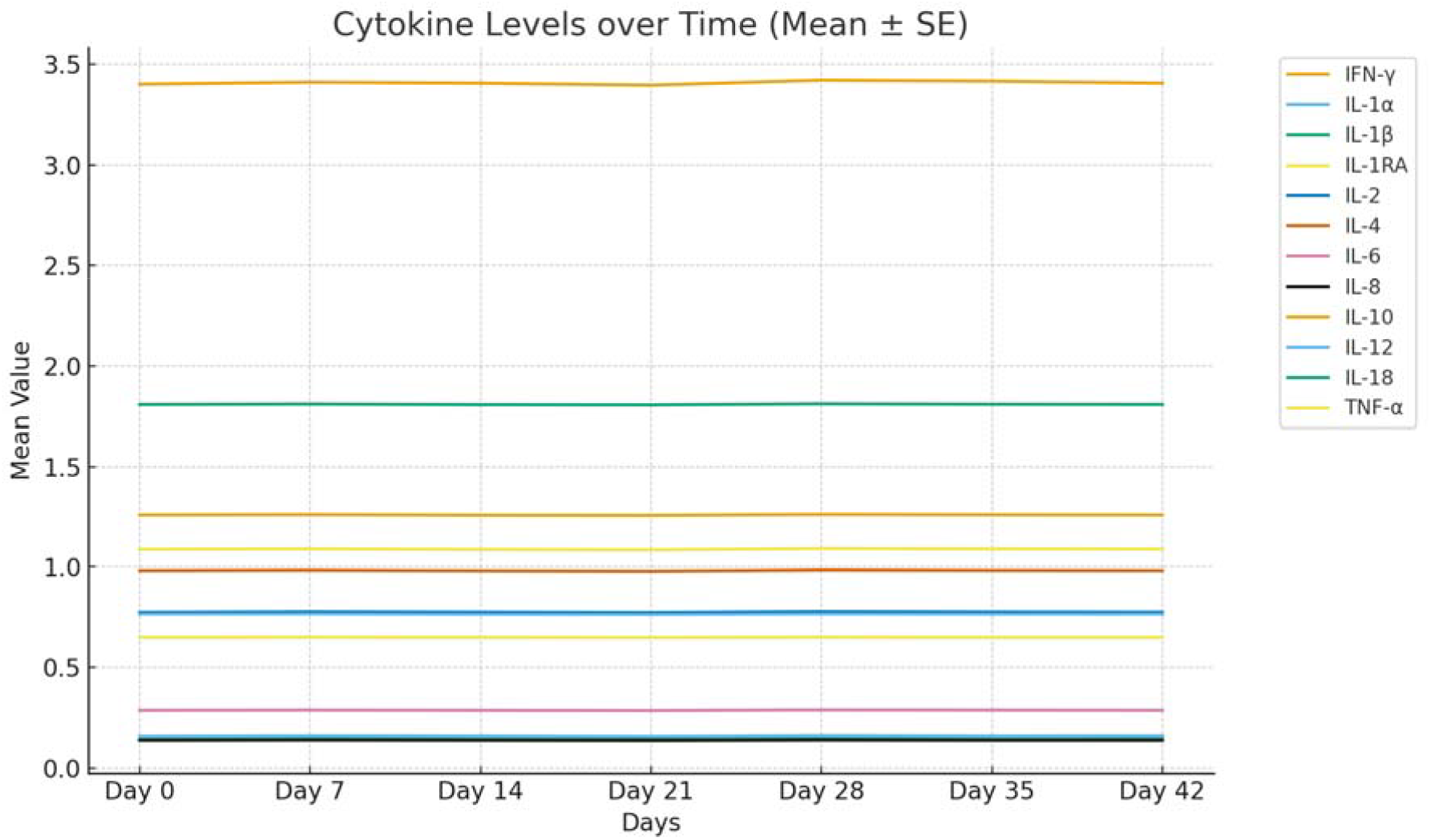
Plasma concentrations of cytokines in non-pregnant pigs.

Further, the pro-inflammatory mediators IL-6, IL-8, IL-12, IL-18, and TNF-α exhibited uniform concentration (Table 2 & Figure 1) with minimal standard error across all sampling intervals. This indicates the absence of any inflammatory or stress-related response, as these cytokines typically rise sharply during immune activation (Ogita and Takai 2008), infection (O’Hayre et al., 2008) or physiological stress. The narrow range of standard errors (SE) across all cytokines reinforces the precision of the measurements and the stability of immune homeostasis throughout the 42-day treatment period. The nearly flat line trends observed in the graphical representation of cytokine concentrations further validate the non-significant temporal variations.

The comparative evaluation of cytokine profiles between pregnant and non-pregnant pigs revealed distinct immunological patterns that underscore the critical role of cytokine modulation during early gestation. In pregnant pigs, there was a significant and progressive increase (p < 0.05) in the concentrations of all analyzed cytokines during early embryonic development stage, suggesting a strong temporal association between cytokine activity and implantation. The marked rise in IFN-γ, IL-1α, IL-1β, and TNF-α indicated their essential involvement in uterine immune modulation, endometrial receptivity and trophoblast invasion processes fundamental for successful implantation. Concurrently, the elevation of IL-1RA demonstrated an adaptive anti-inflammatory response aimed at balancing the pro-inflammatory milieu, thereby preventing excessive inflammation that could lead to early embryonic loss. The significant increase in Th2-type cytokines, particularly IL-4 and IL-10, reflected a well-coordinated shift toward an anti-inflammatory and immunotolerant state at the maternal–fetal interface, facilitating the adhesion of embryo in the uterine body (Robertson et al., 2000). Moreover, the consistent rise in IL-6, IL-8, IL-12, and IL-18 highlighted their synergistic role in vascular remodeling, angiogenesis and immune cell recruitment during implantation (Duc-Goiran et al., 1999) and placental development (Makrigiannakis and Minas,2007). In contrast, non-pregnant pigs exhibited a stable and uniform cytokine profile throughout the 42-day observation period, with no statistically significant fluctuations in any of the measured cytokines. The constant levels of IFN-γ, IL-1 family members, IL-2, IL-4, IL-10, and other pro-inflammatory mediators indicated a steady immunological state without activation of either pro- or anti-inflammatory pathways. This stability confirmed the maintenance of immune homeostasis and the absence of physiological or inflammatory stimuli in non-pregnant animals.

The findings clearly demonstrate that pregnancy in pigs induces a dynamic cytokine network characterized by a balanced interplay between pro-inflammatory and anti-inflammatory mediators. This immunological shift supports endometrial receptivity, implantation and maternal tolerance, processes that are conspicuously absent in non-pregnant animals. Hence, cytokines act as pivotal regulators of cellular communication, immune modulation, and angiogenesis during early pregnancy, reflecting their indispensable role in the successful establishment and maintenance of gestation.

## References

Dey SK, Lim H, Das SK, Reese J, Paria BC, Daikoku T and Wang H. (2004) Molecular cues to implantation. Endocr Rev. Jun;25(3):341–73. doi: 10.1210/er.2003-0020. PMID: 15180948.

Duc-Goiran P, Mignot TM, Bourgeois C, Ferre F. (1999). Embryo-maternal interactions at the implantation site: a delicate equilibrium. Eur J Obstet Gynecol Reprod Biol.;83:85–100.

Engert S, Rieger L, Kapp M, Becker JC, Dietl J and Kammerer U, (2007). Profiling chemokines, cytokines and growth factors in human early pregnancydecidua by protein array. Am J Reprod Immunol; 58: 129–130.

Guzeloglu-Kayisli O, Kayisli UA and Taylor HS. (2009) The role of growth factors and cytokines during implantation: endocrine and paracrine interactions. Semin Reprod Med.Jan;27(1):62–79.

Joyce MM, Burghardt RC, Geisert RD, Burghardt JR, Hooper RN, Ross JW, Ashworth MD, Johnson GA. (2007) Pig conceptuses secrete estrogen and interferons to differentially regulate uterine STAT1 in a temporal and cell type-specific manner. Endocrinology. Sep;148(9):4420–31. doi: 10.1210/en.2007-0505. Epub 2007 May 24. PMID: 17525118.

Kang J and Der SD (2004) Cytokine functions in the formative stages of a lymphocyte’s life. Curr Opin Immunol.; 16:180–190. doi: 10.1016/j.coi.2004.02.002.

Keogh RJ, Harris LK, Freeman A, Baker PN, Aplin JD, Whitley GS, Cartwright JE. (2007) Fetal-derived trophoblast use the apoptoticcytokine tumor necrosis factor-alpha-related apoptosis-inducingligand to induce smooth muscle cell death. Circ Res; 100:834–841.

Kru ssel JS, Biefield P, Polan ML and Simon C. (2003) Regulation of embryonicimplantation. Eur J Obstet Gynecol Reprod Biol; 110 (Suppl. 1):S2–S9.

Makrigiannakis A and Minas V. (2007). Mechanisms of implantation. Reprod Biomed Online. 2007;14:102–109. doi: 10.1016/s1472-6483(10)60771-7.

O’Hayre M, Salanga CL, Handel TM and Allen SJ. (2008) Chemokines and cancer: migration, intracellular signalling and intercellular communication in the microenvironment. Biochem J.; 409:635–649. doi: 10.1042/BJ20071493.

Ogita H and Takai Y. (2008) Cross-talk among integrin, cadherin, and growth factor receptor: roles of nectin and nectin-like molecule. Int Rev Cytol.; 265:1–54. doi: 10.1016/S0074-7696(07)65001-3.

Orsi, N.M. and Tribe, R.M. (2008), Cytokine Networks and the Regulation of Uterine Function in Pregnancy and Parturition. Journal of Neuroendocrinology, 20: 462–469. 10.1111/j.1365-2826.2008.01668.x

Pope WF: 1993. Embryonic Mortality in Swine. In: Embryonic Mortality in Domestic Species. Edited by: Zavy MT, Geisert RD., Boca Raton, CRC Press, 53–78.

Robertson SA, O’Connell A and Ramsey A. (2000). The effect of interleukin-6 deficiency on implantation, fetal development and parturition in mice. Proc Aust Soc Reprod Biol. 2000;31:97.

Ross, J. W., Ashworth, M. D., Hurst, A. G., Malayer, J. R., & Geisert, R. D. (2003). Analysis and characterization of differential gene expression during rapid trophoblastic elongation in the pig using suppression subtractive hybridization. Reprod Biol Endocrinol 1, 23. 10.1186/1477-7827-1-23.

Sands BE, Bank S, Sninsky CA, Robinson M, Katz S, Singleton JW, Miner PB, Safdi MA, Galandiuk S, Hanauer SB, Varilek GW, Buchman AL, Rodgers VD, Salzberg B, Cai B, Loewy J, DeBruin MF, Rogge H, Shapiro M, Schwertschlag US. (1999). Preliminary evaluation of safety and activity of recombinant human interleukin 11 in patients with active Crohn’s disease. Gastroenterology. Jul;117(1):58–64. doi: 10.1016/s0016-5085(99)70550-0. PMID: 10381910.

Sharkey AM and Smith SK. (2003). The endometrium as a cause of implantation failure. Best Pract Res Clin Obstet Gynaecol.; 17:289–307. doi: 10.1016/s1521-6934(02)00130-x.

Sjoblom C, Wikland M and Robertson SA. (1999) Granulocyte-macrophage colony-stimulating factor promotes human blastocyst development in vitro. Hum Reprod; 14: 3069–3076.

Stroband HWJ and Van der Lende T (1990) Embryonic and uterine development during pregnancy. In: Control of Pig Reproduction III J Reprod Fertil Suppl. Edited by: Cole DJA, Foxcroft GR, Weir BJ. Cambridge, UK, 40: 261–277.

